# FAP overexpression induce Epithelial-Mesenchymal Transition (EMT) in oral squamous cell carcinoma by down-regulating DPP9 gene

**DOI:** 10.1101/765743

**Authors:** Qing-qing Wu, Meng Zhao, Guang-zhao Huang, Ze-nan Zheng, Wei-sen Zeng, Xiao-zhi Lv

## Abstract

FAP acts as a tumor promoter via epithelial-mesenchymal transition (EMT) in human oral squamous cell carcinoma (OSCC). The present study was designed to investigate the interaction proteins with FAP and explore the precise mechanism of FAP promoting EMT in OSCC. IP-MS analysis confirmed that DPP9 was an interacting protein of FAP. DPP9 was down-regulated in OSCC tissue samples compared with MNT using immunohistochemistry and quantitative-PCR detection. Lower DPP9 was correlated with unfavorable overall survival of patients with OSCC. Repressing DPP9 accelerates the proliferation of OSCC cells in vitro and in vivo. Mechanistically, overexpression of FAP downregulate the expression of the DPP9 and the effect of FAP on OSCC proliferation, migration, invasion and EMT could be reversed by up-regulated DPP9. Our study suggests that FAP could induce EMT and promote carcinogenesis in oral squamous cell carcinoma by down-regulating DPP9 gene. That will hint different dimension on therapy for patients with OSCC.

## Introduction

Cancer of the oral cavity is one of the most common malignancies and an important cause of morbidity and death(1). OSCC accounts for more than 90% of all oral cancers with the main factors of consumption of tobacco and/or alcohol and chewing areca. In spite of major advances in diagnosis and treatment, the prognosis of OSCC is poor due to invasion, metastasis, and recurrence. Although the oral cavity is easily examined, yet up to 60% of OSCC cases are undiagnosed in the clinical stage. At histopathological level, OSCC is characterized squamous differentiation, nuclear pleomorphisms, invasive growth and metastasis(2). The biomarkers(3) for early diagnosis of OSCC is therefore key to improving patient prognosis and survival rates. FAP is a member of the dipeptidyl peptidase (DPP) family with around 50% homology with DPP4(4). FAP is expressed during development, and in the context of cancer, it is highly expressed and can be a marker of cancer-associated fibroblasts(CAFs), and itself has been demonstrated to have pro-tumorigenic activity(5). Structurally, FAP consists of a 6 amino acid cytoplasmic tail, a single 20 amino acid transmembrane domain, and a 734 amino acid extracellular domain(4) and FAP have both post-proline exopeptidase activity and gelatinase activity(6). FAP plays its role in cancer promotion both by enzymatic effects and non-enzymatic effects. Its dual enzymatic activity gives it a range of putative substrates, and many different types of substrates had been reported(7). Although many studies(7) suggested that FAP can enhance various carcinogenesis process, it is still not clear whether that is based on its enzymatic activity. Emerging(8–10) evidence had suggested FAP’s non-enzymatic role in cancer.

Human cancer FAP expression is reported to correlate to higher tumor grade and worse survival across a worldwide range(11),like cancer on the breast, colon, stomach, etc(12–15). Some studies(15, 16) indicated that FAP can induce EMT. Suppressed FAP expression reduced the adhesion, migration, invasion and metastasis of OSCC cells, and EMT plays a key role in previous studies of these phenotypes(17–20). At the same time, EMT-related marker also changed with reduced FAP, such as high expression of E-cadherin, and low expression of N-cadherin. However, the exact mechanism of FAP in EMT and OSCC carcinogenesis is still unknown. Thus, this study was designed to investigate the possible molecular mechanism of FAP in OSCC.

## Results

### FAP negatively regulate DPP9 in its upstream

The researches currently are limited to its enzymatic activity and its substrates, but the clinical study of the FAP inhibitor Talabostat showed no ideal response. Therefore, it is also important to explore the non-enzymatic activity of FAP. We overexpressed FAP with HIS-tag in SCC9 cell line and conduct IP using anti-HIS antibody to seek the possible interaction proteins with FAP. The lysates from antibody group and IgG group were analyzed by MS, and 14 proteins showed significant difference compared with the IgG group. We focused on DPP9 with 3 filter conditions: subcellular localization, GO function and protein domain^9^ (Fig.1a), and verified it in the antibody group lysate by western blotting (Fig.1b). In SCC9/SCC25 FAP^+^ cell lines, the DPP9 was down-regulated and transiently silencing FAP in SCC15 cell lines, DPP9 expressed lower accordingly. (Fig.1c. *p*<0.05). In order to explore which site on FAP may relate with DPP9, wild type FAP, intracellular segment deletion type FAP (tFAP) and extracellular segment mutation type FAP (mFAP) in pcDNA3.1+ plasmid were transiently transfected into SCC9 cell lines. Then DPP9 antibody was used to execute IP, and the FAP was detected in four groups of lysates, observing no FAP in tFAP group. This suggested that intracellular part of FAP may be the cooperating site with DPP9. (Fig.1d) To confirm FAP will not be regulated by DPP9, when transfecting siRNA to silence DPP9, we detected FAP expression in both SCC9 and SCC25 groups and observe FAP no difference (Fig.3a).

**Figure 1.**
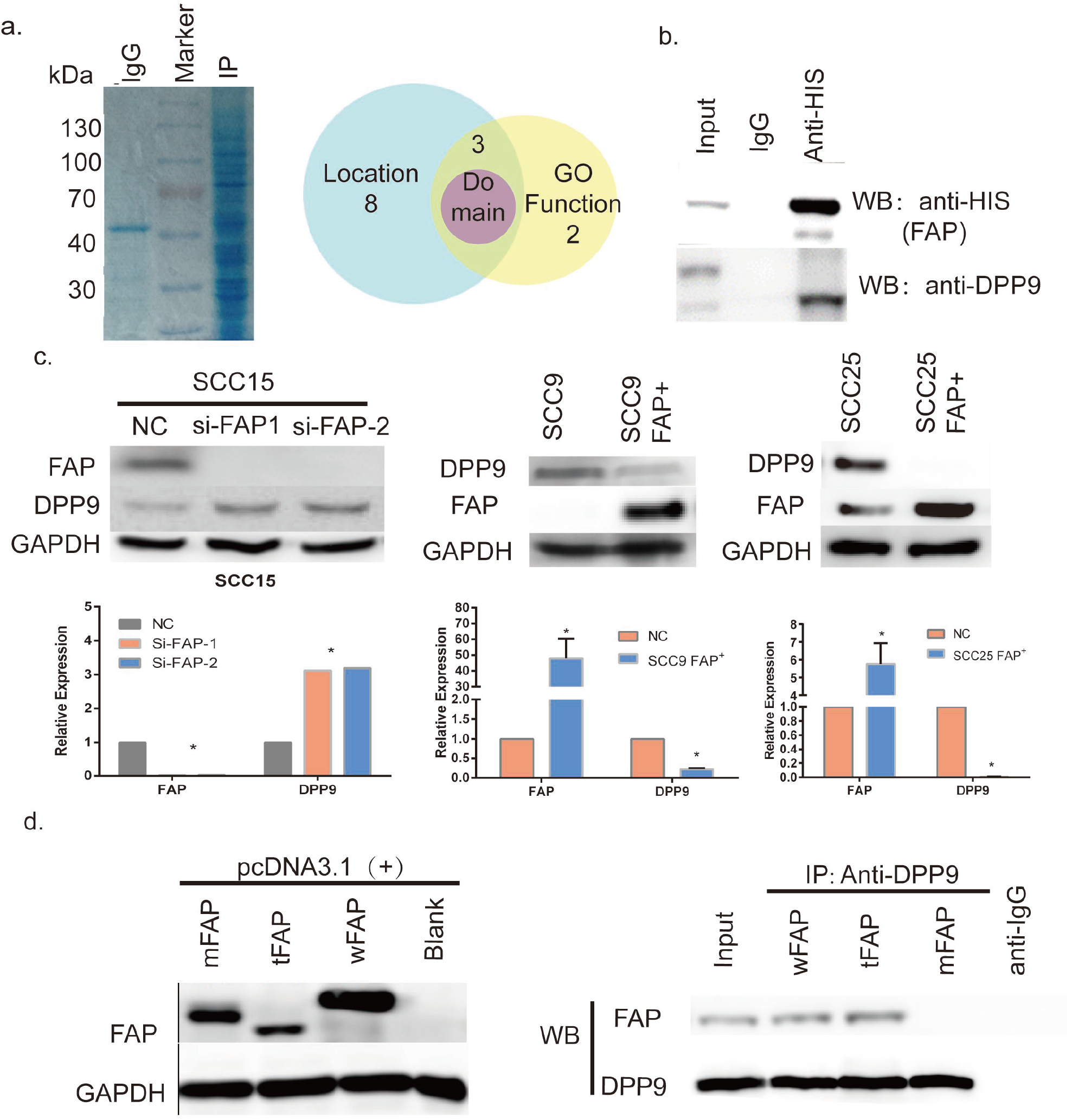
Negative correlation between FAP and DPP9. Coomassie blue staining shows IP lysate with His-antibody. Venn diagram shows the filter condition of 14 peptides from IP-MS. **(b)** Verify DPP9 in IP lysate using DPP9 antibody. **(c)** In FAP-low-expression SCC9 and SCC25, overexpressing FAP and reduced DPP9 can be detected. And in FAP-high-expression SCC15, knockdown FAP shows higher DPP9. All data are represented as mean ± SD; *p < 0.05. **(d)** Using pcDNA-wFAP/mFAP/tFAP transfecting SCC9 and IP by DPP9 antibody show no blot in tFAP group compared with wFAP/mFAP group.

### DPP9 is downregulated in OSCC

Through analysis of DPP9 expression from TCGA head and neck cancer patients in Oncomine, the log^2^ copy number unit of DPP9 was downregulated in head and neck carcinoma samples (290 cases) compared with head and neck part (74 cases) and blood samples (264 cases) (Fig.2a). Three randomized-picked paired OSCC tumor tissues (TUM) and the MNT were analyzed and western blotting revealed that DPP9 protein were markedly low expression in the tumor tissues compared to the MNT. (Fig.2c). DPP9 mRNA expression in Tum was relatively lower in MNT. (*p*< 0.0001, n=90, Fig.2d). Also, protein expression levels of DPP9 were measured in samples of 118 pairs archived paraffin-embedded TUM and MNT using IHC (*p*< 0.005, n=45) (Fig.2e). Taken together, these results strongly indicate that DPP9 is downregulated in human OSCC.

**Fig.2.**
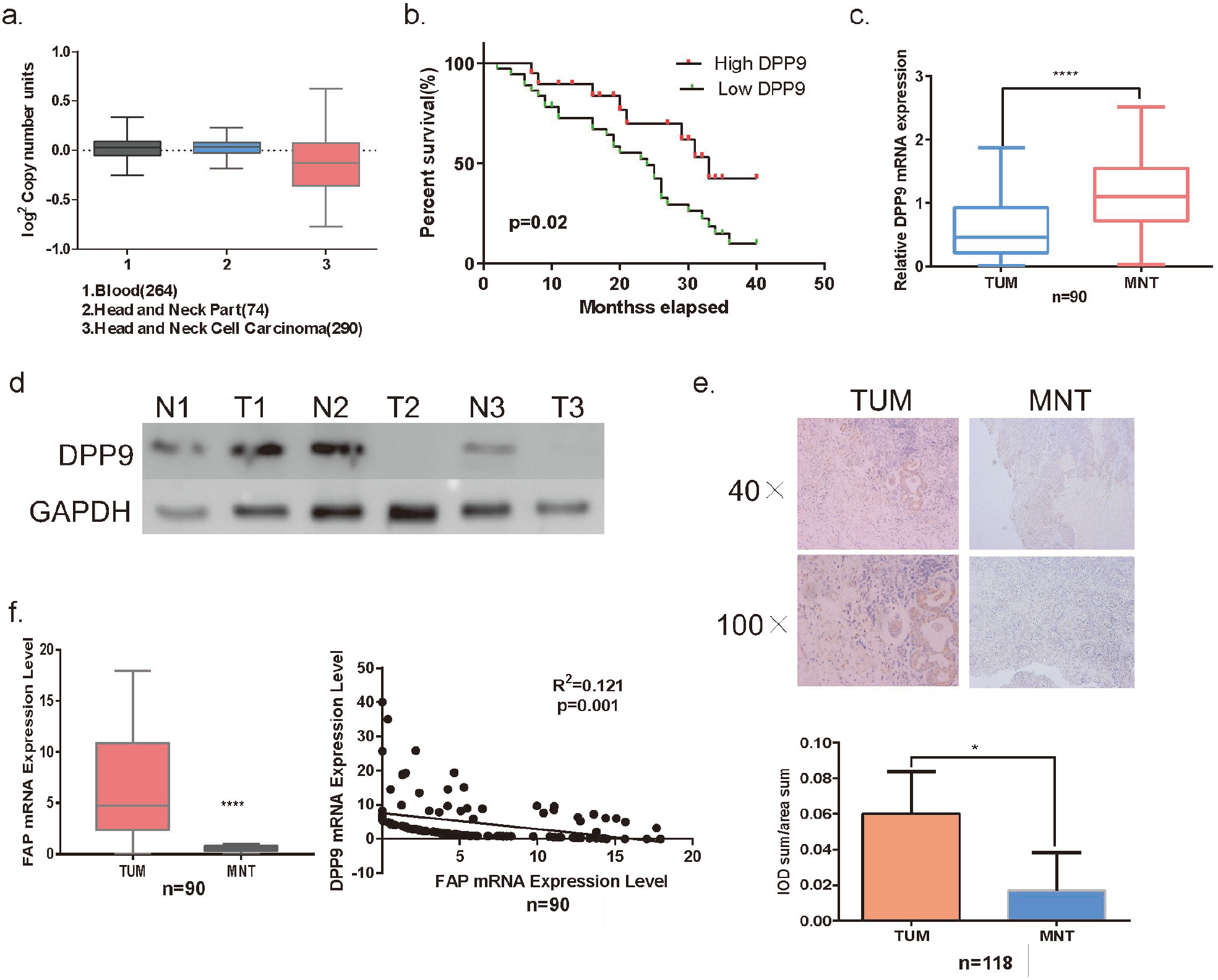
In Oncomine database (The Cancer Genome Atlas - Head and Neck Squamous Cell Carcinoma DNA Copy Number Data), HNSCC shows lower DPP9 copy number units compared with that in the blood sample and the Head and Neck part. **(b)** Survival curves (Kaplan–Meier plots) show low-DPP9 is related to a lower survival rate(*p*=0.02). **(c)** mRNA level of DPP9 in 90 pairs clinical samples. **(d)** Three random pairs of samples detecting the DPP9 protein expression. **(e)** By IHC and statistical analysis show DPP9 protein level is less in TUM compared with MNT(\*\*\*\**p<*0.0001). **(f)** mRNA level of DPP9 in 90 pairs clinical samples and egression analysis showed that DPP9 is negatively associated with FAP in OSCC tissues.

### Decreased expression of DPP9 is unfavorable for OSCC prognosis

To explore the prognostic value of DPP9 expression for OSCC, the follow-up data of 118 OSCC patients for up to 40months were used to assess the value of DPP9 for predicting patient survival in OSCC patients. These samples were stained using a DPP9 antibody and scored using a standard method (summarized in Table 1). Compared to the adjacent non-tumor OSCC in which DPP9 was strongly detected, DPP9 was undetectable or found to be only expressed at low levels, DPP9 was low expressed in OSCC specimens (Fig.2c). We analyzed the association between DPP9 and the clinicopathological features of OSCC. As showed in Table 1, strong associations were observed between DPP9 expression and clinical stage (*p*=0.036), T classification (*p=*0.017), N classification(*p*=0.041). However, the expression of DPP9 was not associated with age, gender and Lymphatic metastasis. Kaplan–Meier survival analysis revealed a correlation between DPP9 expression level and overall survival times (*p*=0.02, Fig.2b). These results indicate a significant correlation of the expression of DPP9 with the prognosis of OSCC.

**Table1:**
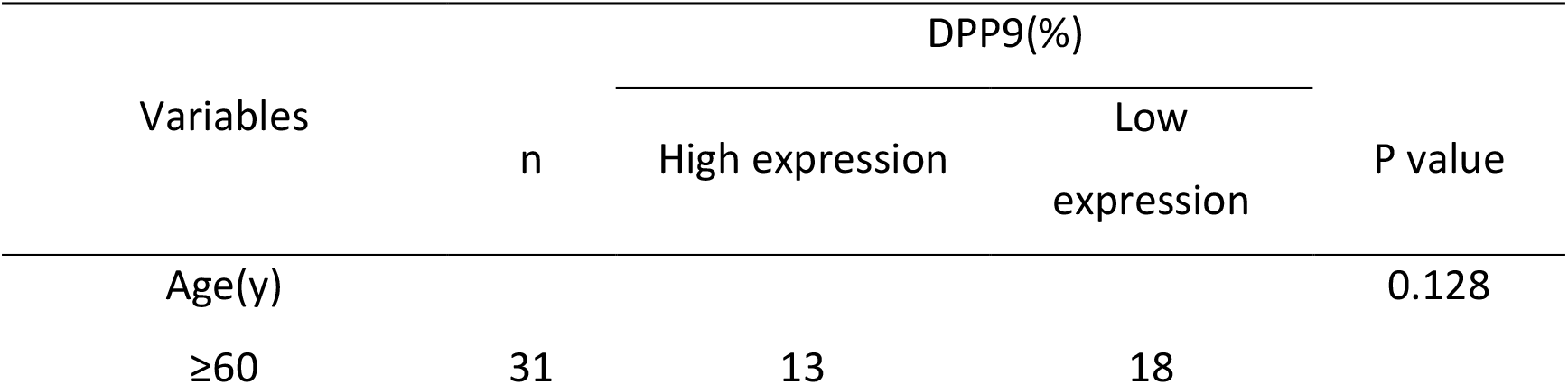
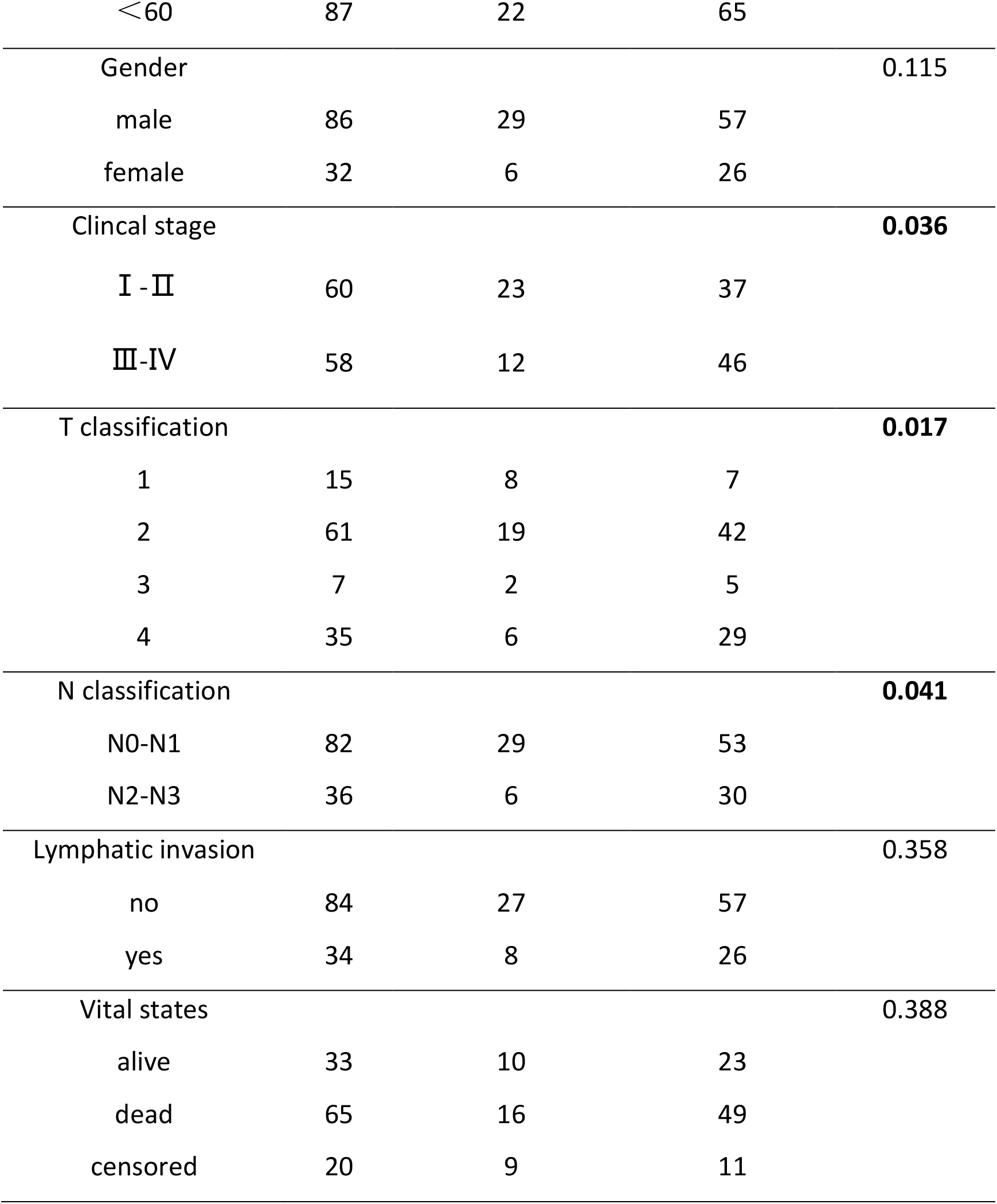
Clinicopathological characteristics of patient samples and expression of DPP9 in OSCC

| Variables | n | DPP9(%) |  | P value |
| --- | --- | --- | --- | --- |
|  |  | High expression | Low expression |  |
| Age(y) |  |  |  | 0.128 |
| ≥60 | 31 | 13 | 18 |  |
| <60 | 87 | 22 | 65 |  |
| Gender |  |  |  | 0.115 |
| male | 86 | 29 | 57 |  |
| female | 32 | 6 | 26 |  |
| Clinical stage |  |  |  | **0.036** |
| I - II | 60 | 23 | 37 |  |
| III-IV | 58 | 12 | 46 |  |
| T classification |  |  |  | **0.017** |
| 1 | 15 | 8 | 7 |  |
| 2 | 61 | 19 | 42 |  |
| 3 | 7 | 2 | 5 |  |
| 4 | 35 | 6 | 29 |  |
| N classification |  |  |  | **0.041** |
| N0-N1 | 82 | 29 | 53 |  |
| N2-N3 | 36 | 6 | 30 |  |
| Lymphatic invasion |  |  |  | 0.358 |
| no | 84 | 27 | 57 |  |
| yes | 34 | 8 | 26 |  |
| Vital states |  |  |  | 0.388 |
| alive | 33 | 10 | 23 |  |
| dead | 65 | 16 | 49 |  |
| censored | 20 | 9 | 11 |  |

### DPP9 modulates cell growth of OSCC cells

To evaluate the functional significance of DPP9 on cancer cell proliferation, si-RNA was transfected into SCC9 and SCC25 cell lines to specifically knockdown the expression of DPP9. The effect of siRNA transfection on the expression of DPP9 was confirmed by western blot analysis (Fig.3a, \*\**p<*0.01). DPP9 knockdown in SCC9 and SCC25 cells promoted cell growth in vitro. Colony formation assay showed that suppressing DPP9 significantly stimulated cell proliferation (Fig.3b). The growth curves determined by CCK-8 assay showed that suppression DPP9 significantly stimulated cell viability in comparison with si-con cells (Fig.3c). Taken together, these results suggested that DPP9 significantly inhibited cell growth in vitro.

**Fig.3.**
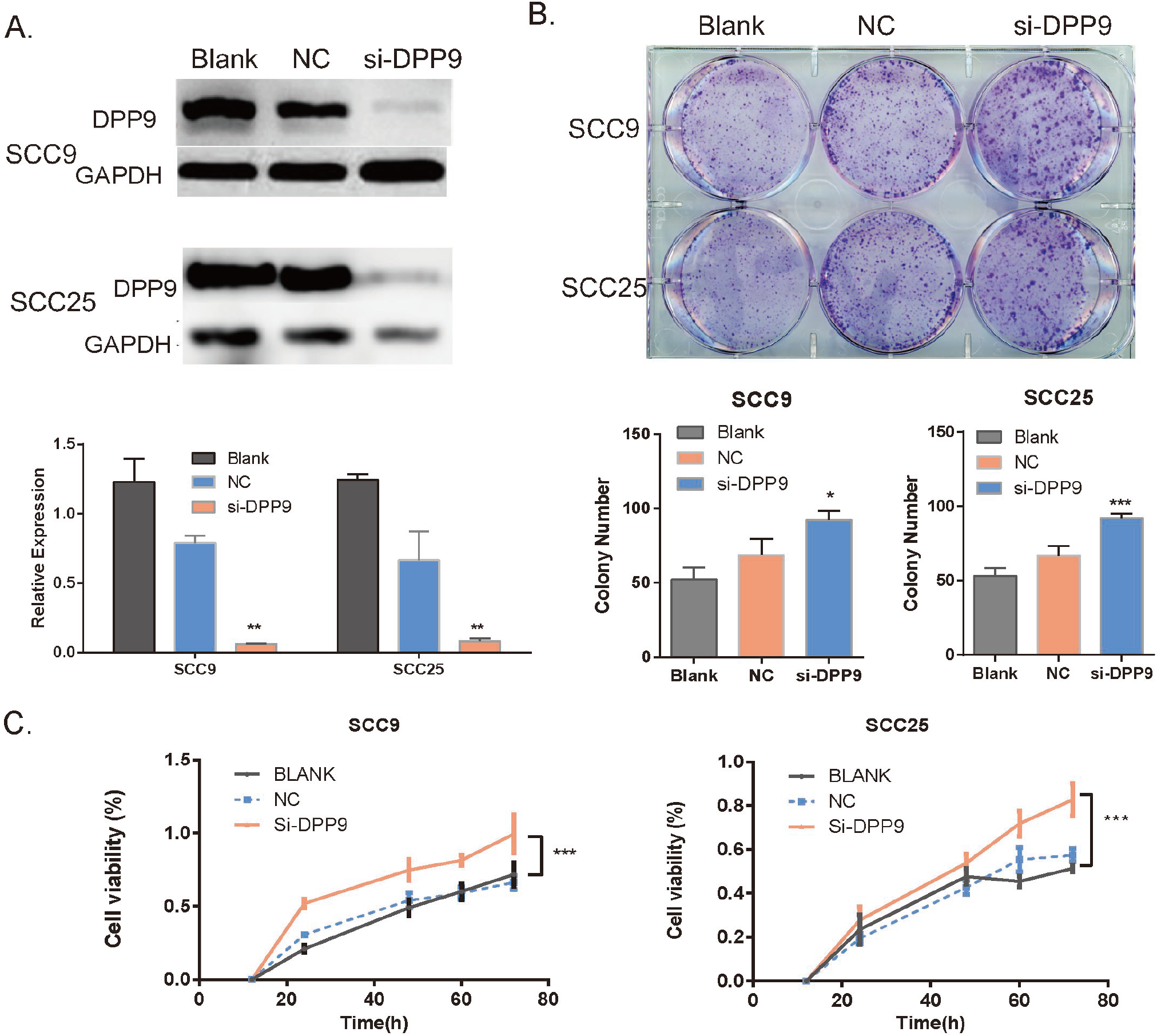
Knockdown of DPP9 causes an anti-tumor effect on OSCC cells. **(a)** Western blot analysis of DPP9 following DPP9 knockdown for 72h. GAPDH was used as an internal control. **(b)** Colony formation assay after knockdown of DPP9 in SCC9 and SCC25cells for 10days (*Up*). The mean number of colonies for each well was determined from three independent assays (*Down*). **(c)** Growth rates of SCC9 and SCC25 cells measured by CCK-8 assay after DPP9 knockdown. All data are presented as mean ± SD; *p < 0.05; **p < 0.01; ***p < 0.001 versus control group.

### Repression of DPP9 promotes cell migration and invasion in vitro

To investigate the effects of DPP9 on cancer cell metastatic ability, repression of DPP9 in SCC9 and SCC25 cells was conducted to examine the cellular function of invasion and migration. In a wound-healing assay, there was a marked acceleration in the silencing group at the edges of the scratch wound of SCC9 and SCC25 cells (Fig.3c) and Quantitative analysis at 24h confirmed a significant promotion. In addition, Matrigel invasion assays showed SCC9 and SCC25 cells with downregulated DPP9 were much more invasive than controls and the similar tendency in migration assays (Fig.4a, \**p<*0.05, **p<0.01, \*\*\**p<*0.005).

**Fig.4.**
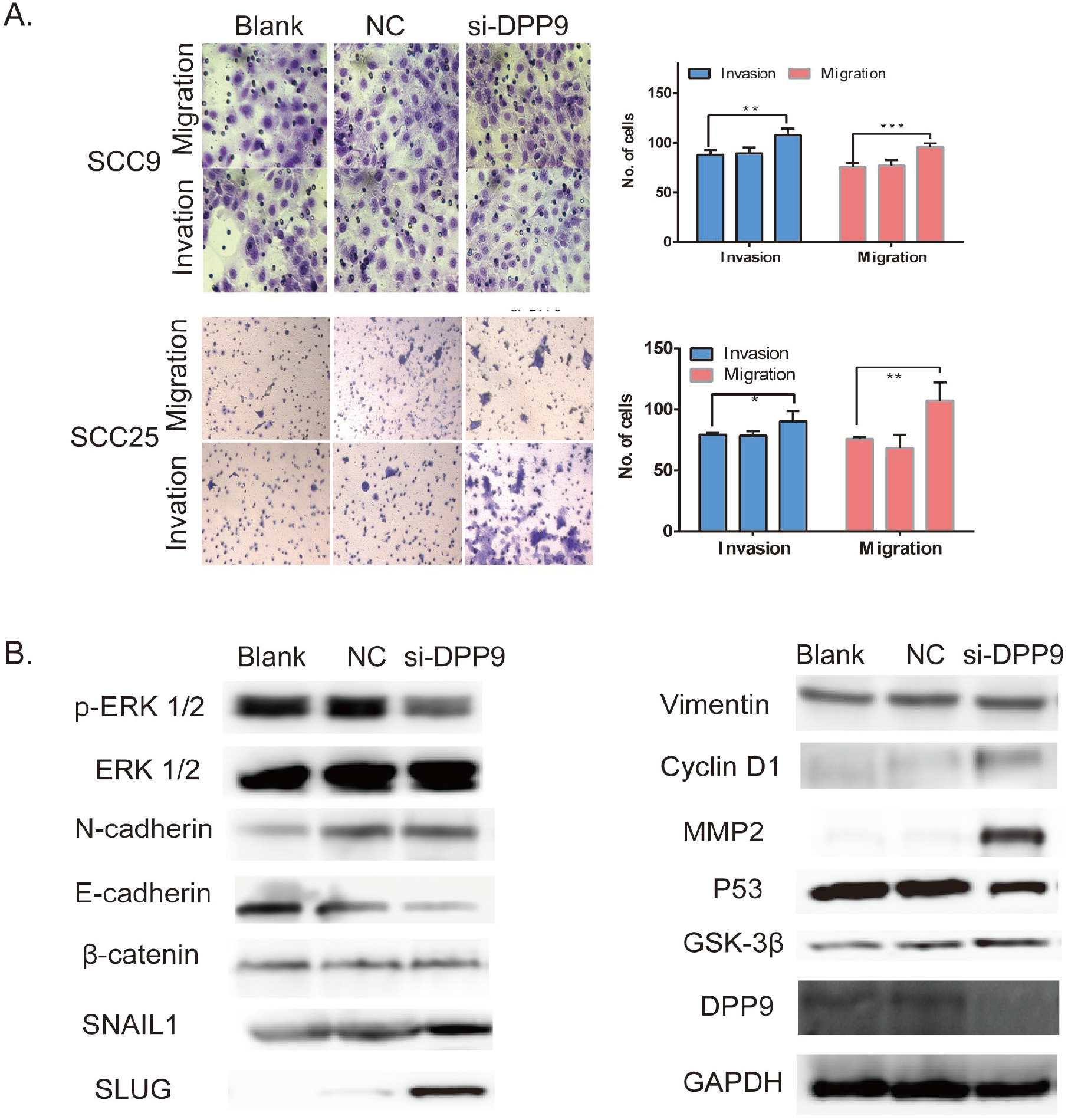
Depression DPP9 promoted cell migration/invasion and induced EMT in vitro. **(a)** Transwell migration and invasion assay after DPP9 knockdown for 48h. All data are presented as mean ± SD; \**p* < 0.05; \*\**p* < 0.01; \*\*\**p* < 0.001. **(b)** Western blot analysis of p-ERK1/2, ERK, EMT-associated proteins and P53, DPP9, Cyclin D1 following knockdown for 48h. GAPDH was used as an internal control.

### DPP9 modulates the expression of multiple genes involved in the EMT in OSCC

Still, use si-DPP9 into SCC9 and SCC25, EMT-associated markers were detected, where the protein level of E-cadherin was decreased while the N-cadherin, Vimentin, SNAIL, was markedly increased in compared with control cells in vitro (Fig.4b). And the phosphorylated ERK1/2 is significantly decreased compared with the control group with similar pan-ERK1/2 expressions. And the proliferation associated protein Cyclin D1, BCL-2 is also detected and shown higher expression.

### DPP9 modulates cell growth of OSCC cells in vivo

To assess the effect of DPP9 of OSCC growth in vivo, DPP9-depleted SCC9 cells, or control cells were injected into nude mice subcutaneously, and then monitored tumor growth. Tumor generation speed of si-DPP9 cells was significantly faster than in control cells, and final volume/weight (20 days) of si-DPP9 group is significantly larger than the SCC9 group(Fig.5a). And the EMT-associated protein in si-DPP9 group change similarly with treated cells in vitro compared with the untreated group by WB (Fig.5b). All tumors were taken into HE staining and IHC staining using DPP9, CD44, Ki67 (Fig.5c).

**Fig.5.**
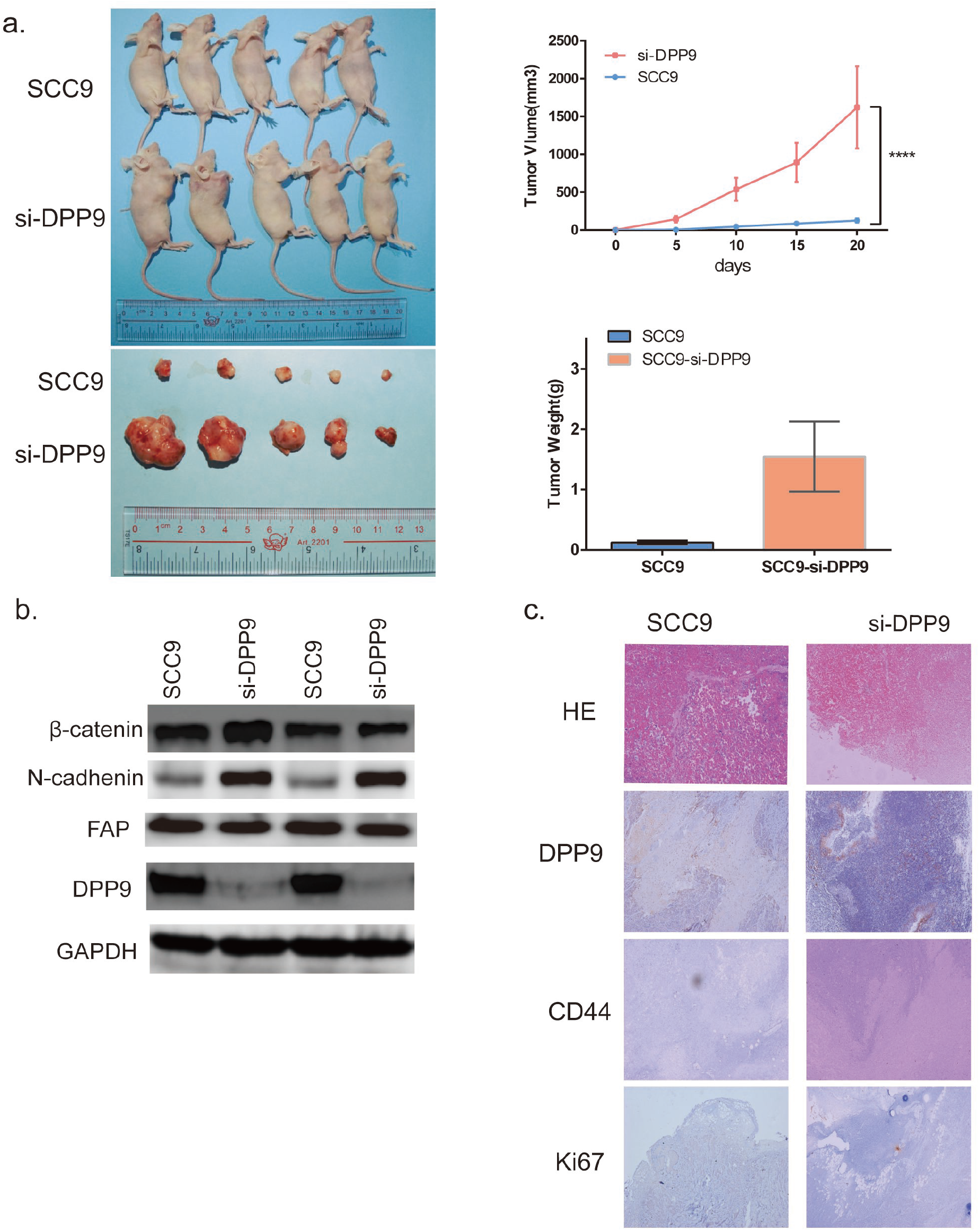
DPP9 modulates cell growth of OSCC cells in vivo. **(a)** Tumor formation for 20days after injecting cells to BALBc/nude right armpit. The weight of the tumors and volume change were observed every five days. And the mean weight of tumors for each group was showed in bar graph. **(b)** Western blot analysis of DPP9, FAP, EMT-associated proteins in two tumor tissues each group. **(c)** IHC of CD44, Ki-67 and DPP9 verification and HE staining of each group.

### Cell proliferation and migrative/invasive promotion caused by overexpressing FAP is associated with the DPP9 in OSCC cells

To further delineate the role of DPP9 in the proliferation and migration/invasive regulated by FAP, the expression of DPP9 was restored by pCMV-DPP9 plasmid in FAP-overexpressing SCC9 cells (Fig.6a). As showed in Figure 6, results of the CCK8 assay and colony formation assay demonstrated that the DPP9 plasmid decreased the cell proliferation and tumorigenesis (Fig.6b,6e). Consistently, cells numbers of DPP9 plasmid treated cells of migration and invasion was reduced (Fig.6d). Also, the adhesion test showed further reduction of adhesion ability of the treated cells (Fig.6c). And Western blotting showed reversed expression of, SNAIL1, Vimentin, E-cadherin and N-cadherin, and no significant change of Cyclin D1, β-catenin and BCL-2(Fig.6f). Taken together, these findings suggest that the observed regulation of proliferation, tumorigenesis caused by overexpressing FAP is associated with the DPP9 in OSCC.

**Fig 6.**
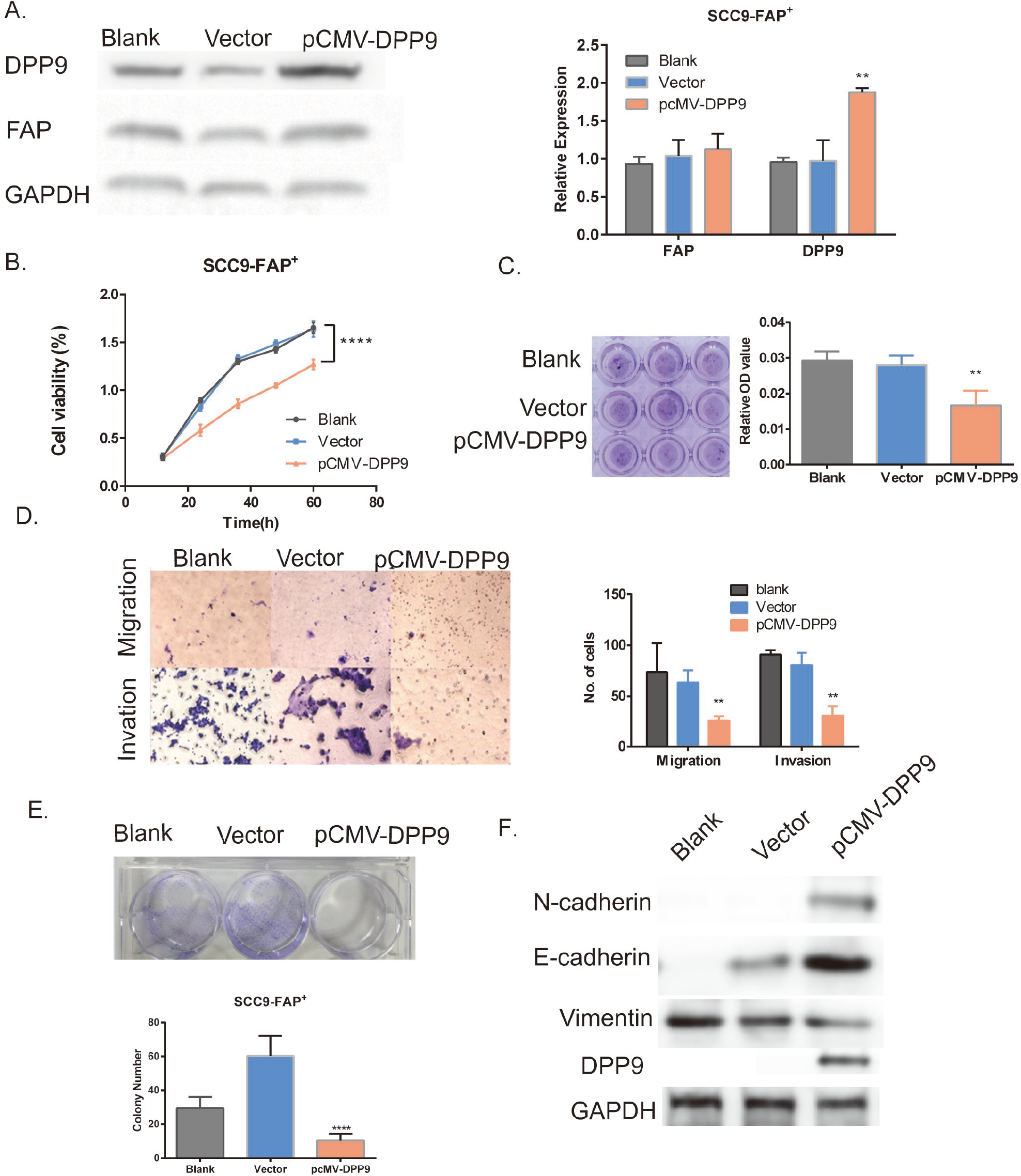
Cell proliferation and migrative/invasive promotion caused by overexpressing FAP is associated with the DPP9 in OSCC cells. **(a)** Verification of DPP9 overexpression in SCC9-FAP^+^ cells by Western Blotting. **(b)** CCK-8 shows decreased growth rate of treated cells. **(c)** Transwell migration and invasion assay after DPP9 overexpressing 48h. **(d)** *Up:* Colony formation assay after DPP9 plasmid transfecting for 10days. *Down:* The mean number of colonies for each well was determined from three independent assays. **(e)** Adhesion test showed further reduction of cell adhere ability in the treated group. **(f)** Western blot analysis of N-cadherin, E-cadherin and Vimentin level in SCC9-FAP^+^ cells after treatment with pCMV-DPP9 for 48h. All data are presented as mean ± SD; \**p* < 0.05; \*\**p* < 0.01; \*\*\**p* < 0.001 versus control (Blank, Vector)

## Discussion

FAP has been considered as a potential immunotherapeutic target(21), and it plays a key role in cancer promotion both by enzymatic effects and non-enzymatic effects. Its enzymatic effects attract more attention so far. Its substrates are partially shared with DPP4(22),(23). However, Talabostat, the first inhibitor of enzymatic activity of FAP, showed no significant response in clinical trials(24–26). Emerging evidence supports the argument that FAP might affect tumorigenicity independent of its enzymatic activity. Expressing FAP and mutation FAP, both displayed enhanced cancer phenotypes with no significant difference(8). Chung suggested FAP leads to migration and its peptidase activity is not essential for such migration(10). Yang et al.(9) Inhibited FAP with Talabostat does not induce levels of its downstream CCL-2 secretion. Knopf et al.(27) suggested FAP direct bound to erlin-2, stomatin, and caveolin-1. However, it is not clear whether FAP binds to erlin-2, stomatin and caveolin-1 molecules through enzyme activity pattern or non-enzyme activity pattern. FAP may bind to erlin-2 through a non-enzymatic pattern for erlin-2 locating on the endoplasmic reticulum. Therefore, we are committed to exploring the closest downstream effectors on/inside the cell that may mediate the regulation of proliferation and metastasis of FAP, and in 14 proteins analyzed by IP-MS. 2 of 14 proteins are located on the cell membrane and the other 12 are distributed separately on nucleus, cytoplasm, mitochondrion, cytosol, cytoskeleton. We finally focus on DPP9 because of its similar location, likely domain and analogous GO function compared to FAP.

DPP9 belongs to the DPP gene family(28), locates in cell cytosol, expresses ubiquitously in human tissues and mainly enriched in lymphocytes and epithelial cells(29). DPP9 and DPP4 share a high sequence homology, and possess a very similar tertiary structure and functional activity. Not like DPP4, the intracellular functions of DPP9 in human cancer are still unclear. DPP9 has two distinct biological effects on transformed cells in various types of cancer. On the one hand, Lu et al.(30) found DPP9 may act as a survival factor for cells from the Ewing’s sarcoma family of tumors cells. Yao(31) reported that DPP9 overexpression can inhibit PI3K/AKT signaling, attenuate cell proliferation and promoting apoptosis in human hepatoma cells(32). On the other hand, DPP9 expression is significantly increased in testicular cancer(29). Thence the pro-or anti-tumor activity of DPP9 may depend on the cell type and the molecular context within the tumor microenvironment. The functional role of DPP9 in OSCC remains to be elucidated.

In our study, we verified that DPP9 is significantly downregulated in OSCC samples both from TCGA database and in our sample base and the lower DPP9 expression is associated with worse prognosis, including clinical stage, T, N classification and lymph node status. And survival analysis showed that downregulated DPP9 was an independent prognostic factor for poor 3-year overall survival in OSCC patients. Loss-of-function experiments indicated that DPP9 knockdown fostered the colonies forming, cell proliferation speed, migration and invasion of OSCC cells. All these results support an anti-cancer role for DPP9 in OSCC. To the best of our knowledge, this is the first study to explore the clinical prognostic value of DPP9 in OSCC.

EMT has been confirmed to play a significant role in promoting metastasis in epithelium-derived carcinoma. Accumulating evidence has shown that EMT confers adhesion, migration, invasion and metastasis capacity, stemness and multidrug resistance in tumor cells(17–20). When repressing DPP9, the protein level of E-cadherin, which is a hallmark of EMT(19, 33, 34), was decreased while the N-cadherin, SLUG, SNAIL1 was markedly increased in compared with control cells in vitro. And the Vimentin and β-catenin expression cannot be observed changed. We also detect the biomarkers of proliferation and metastasis that may be mediated by DPP9. The expression of MMP2 and Cyclin D1 enhanced while P53 and phosphate-ERK1/2 decreased and GSK-3β show no difference. Thus, our data and results conclude that the underlying mechanism of DPP9 towards OSCC maybe the EMT properties regulation. Currently, there is not much research on whether DPP9 affects EMT, while Tang.(35) found upregulated DPP9 promotes tumorgenicity and EMT in NSCLC. This difference may be because NSCLC samples in Tang’s study included adenocarcinoma and squamous cell carcinoma, which may have a certain impact on the study.

It can be observed the reversed phenotype of tumor growth, migration and invasion when enforcing DPP9 expression in SCC9-FAP^+^ cells. And our conditioned-IP revealed that the intracellular part of FAP might be the possible site interact with DPP9. We reach the conclusion that FAP facilitates cell phenotypes and EMT by down-regulating DPP9. This results innovatively indicate that FAP may induce pro-tumorigenic effects in a non-enzymatic manner. The extracellular segment containing enzyme active site may not be the only part to promote tumor.

## Materials and methods

### Cell culture, tissue collection, and Ethics Statement

OSCC cell lines SCC9, SCC25, SCC15 were purchased from ATCC and maintained in DMEM/F12 supplemented with 10% newborn calf serum (NBCS) (Gibco Company, USA). In all, 118 OSCC specimens and matched normal tissues (MNT) were obtained at the time of diagnosis before any therapy from Nanfang Hospital of Southern Medical University, Guangzhou, from 2015 to 2018. In 118 cases, there were 86 males and 32 females. For the use of these clinical materials for research purposes, prior written informed consents from all the patients and approval from the Ethics Committees of Nanfang Hospital of Guangdong Province was obtained (NO: NFEC-2018-027). All specimens had confirmed the pathological diagnosis and were staged according to the 2009 UICC-TNM Classification of Malignant Tumors.

### Transient transfection with siRNAs for FAP and DPP9

siiRNA for FAP and DPP9 was designed and synthesized by Genepharma (GenePharma Inc., Suzhou, PR China). Twenty-four hours before transfection, SCC9 cells were plated onto a 6-well plate (Jet Bio-Filtration Co., Ltd, Guangzhou, PR China) at a 30–50% confluence. They were then transfected into cells using Lipofectamine3000 Transfection Reagent (Thermos Fishers Co, Ltd., USA) according to the manufacturer’s protocol. Cells were collected after 48–72 h for the further experiments.

### RNA isolation, reverse transcription, and qRT-PCR

Total RNA was extracted from the cells using Trizol (Takara, Shiga, Japan). Reverse transcription and qPCR were performed in accordance with the manufacturer’s instructions (Vazyme Biotech Co., Ltd, Nanjing). The PCR for each gene was repeated three times. GAPDH was used as an internal control to normalize FAP and DPP9 expression. Differential expression of FAP was calculated using the method.

### Plasmid Construction

PFU enzyme (Thermo Fisher, Inc; USA) was used for the PCR program. Using the cDNA from the OSCC samples as template, fragment wild type FAP (wFAP) with HIS-tag was cloned out with primer A and primer B.

Taking wFAP as template, for tFAP (intracellular site deleted), we use primer C and primer B to clone out fragment tFAP. And for mFAP (site 624-704 deleted), we had overlap primer D and primer E. After using primer A and D, primer B and C respectively to clone out fragments F1 and F2, both fragments were added together as templates and cloned out mFAP with primer A and primer E. These three fragments (wFAP, tFAP, mFAP) was cloned into plasmid pcDNA3.1(+) using BamHⅠand EcoRⅠ. Plasmids was transfected into cells with Lipofectamine3000 Transfection Reagent likely mentioned before. Using the cDNA and primer F/G to clone fragment DPP9 and set it into plasmid pCMV3-Flag with KpnⅠand XbaⅠ.

### Western blotting

Western blotting was performed using a SDS-PAGE Electrophoresis System according to the previous descriptions with rabbit polyclonal anti-FAP antibody (1: 1000; Santa Cruz Biotechnology Inc., Seattle, USA), mouse monoclonal anti-DPPR2 antibody (1:2000; Santa Cruz Biotechnology Inc., Seattle, USA). Another rabbit polyclonal antibodies contain HIS-tag, GAPDH, E-cadherin, N-cadherin, Vimentin, MMP2, P53, SNAIL, SLUG (1:2000; Proteintech Inc., USA) and pan-ERK1/2 and phosate-ERK1/2(1:2000; CST, USA).

### IP assay and MS assay

The total protein was extracted in SCC9-FAP and IP was performed using a Protein A/G Magnetic Beads (Bimake, USA). Briefly, SCC9-FAP^+^ cells were washed with phosphate-buffered saline (PBS), lysed in cold IP lysate buffer, and then centrifuged. Next, the cell lysates were immunoprecipitated with the monoclonal anti-his-tag antibody (Cell Signaling Technology, Inc.; USA) and incubated overnight at 4°C. SDS-PAGE was then conducted to examine the protein levels. The lysate including his-FAP-interacting peptides were collected and submit to a MS facility (Thermo Fisher, Inc.; USA) for analysis. The experiments were repeated three times and the detected peptides were intersected.

### Immunohistochemistry (IHC) assay

Examination of DPP9 expression in samples of OSCC and its control tissues by IHC was processed with the DPP9 antibody (1: 50, Abcam Inc., USA). The stained tissue sections were reviewed and scored independently by two pathologists blinded to the clinical parameters. The staining score standard has also been described. For statistical analysis of DPP9 expression in noncancerous tissues against OSCC tissues, staining scores of 0–5 and 6–10 were respectively considered to be a low and high expression.

### CCK-8 assay

Cell Counting Kit-8 assay was used to evaluate the rate of in vitro cell proliferation. For siDPP9 cells, they were seeded in 96-well plates at a density of 1000 cells/well and respectively incubated for 12, 24, 48, 70 hours. 10μl of CCK-8 (DOJINDO LABORATORIES Ltd., China) was added to each well and incubated for 4 h. The absorbance value (OD) of each well was measured at 450nm. Experiments were carried out three times.

### Colony formation assay

Placing 1,0000 treated cells per well in 6-well plates and after their attachment, there is no contact between cells. Incubate the cells in a CO_2_ incubator at 37°C for 10-15days until cells have formed colonies with substantial good size (50 cells per colony). Remove medium and rinse cells with PBS 2 times, and add 1ml methanol per well at room temperature (RT) for 5 min. Remove methanol and add 0.5% crystal violet carefully and incubate at RT for 2h. Remove crystal violet and immerse the plates in tap water to rinse off crystal violet. Air-dry the plates at RT for a day. Scan the plates into image and count number of colonies, then calculate plating efficiency(PE) by the quotation PE=no. of colonies formed/no. of cells seeded×100%.

### Invasion and migration assays

For the invasion assay, cells were seeded in 100 ml DMEM/F12 media on the top of polyethylene terephthalate (PET) membranes coated with Matrigel TM (1.5 mg/ml, BD Biosciences Inc.) within transwell cell culture inserts (24-well inserts, 8 mm pore size; Corning Life Sciences, Corning, NY, USA). The bottom chamber was filled with 600 ml of DMEM/F12 media containing 20% FBS. The cells were incubated for 12 h at 37°C with 5% CO2. Subsequently, the cells were fixed in 2.5% (v/v) glutaraldehyde and stained with 0.1% crystal violet. For the migration assay, the same of the invasion assay without Matrigel TM. Both cells on the membranes bottom were visualized under a microscope (Zeiss Ltd., China) and quantified by counting the number of cells in three randomly chosen fields at 200-fold magnification.

### In vivo tumor growth assay

BALB/c-nude mice (4 weeks old, 18-20g) were purchased from The Laboratory Animal Centre, Southern Medical University. The Institutional Animal Care and Use Committee of Southern Medical University approved all experimental procedures. Cells were harvested by trypsinization, washed twice with cold serum-free medium, and resuspended with serum-free medium. To evaluate cancer growth in vivo, 2 × 10^6^ treated cells were independently injected subcutaneously into the left axilla in 5 nude mice each group. Every five days, the length (L) and width (W) of tumors were measured using calipers, and their volumes were calculated using the equation (L ×W^2^)/2. On day 20th, the animals were euthanized, and the tumors were excised, weighed, serial sliced and stained with hematoxylin and eosin (HE).

### Statistical analysis

SPSS 24.0 software (SPSS Inc., Chicago, IL, USA) and GraphPad software (GraphPad Software, Inc., La Jolla, CA, USA) was used to analyze all data for statistical significance. Two-tailed Student’s t-test was used for comparisons of two independent groups. One-way ANOVA was used to determine differences between groups for all in vitro analyses. A *p*<0.05 was considered statistically significant.

## Conclusion

Overall, our study suggests that FAP induces EMT by down-regulating DPP9 gene in OSCC. DPP9 may play a potential tumor depression role in OSCC.DPP9 overexpression was able to reverse tumorigenesis and metastasis caused by FAP in OSCC cells. These could be a vigorous therapeutic strategy for future OSCC treatment, but the specific mechanism and functional sites still need to be further explored.

## Abbreviations

FAP: fibroblast activation protein
OSCC: Oral squamous cell carcinoma
IP: Immunoprecipitation
MS: Mass spectrometry
qPCR: Quantitative real-time PCR
EMT: Epithelial-mesenchymal transition

## Acknowledgments

QQW carried out her thesis research under the auspices of the NanFang Hosipital, Southern Medical University and Department of Cell Biology, School of Basic Medical Science, Southern Medical University, Guangzhou, China.

## Ethics approval and consent to participate

The study protocol was approved by the Ethics Committees of Nanfang Hospital of Guangdong Province (NO: NFEC-2018-027).

## Availability of data and materials

The datasets used and analyzed during the current study are available from the corresponding author on reasonable request.

## Consent for publication

Not applicable.

